# Retinoic acid accelerates the specification of enteric neural progenitors from *in vitro*-derived neural crest

**DOI:** 10.1101/819748

**Authors:** Thomas J.R Frith, Antigoni Gogolou, James O.S Hackland, Ivana Barbaric, Nikhil Thapar, Alan J. Burns, Peter W Andrews, Anestis Tsakiridis, Conor J. McCann

## Abstract

The enteric nervous system (ENS) is derived primarily from the vagal neural crest, a migratory multipotent cell population emerging from the dorsal neural tube between somites 1-7. Defects in the development and function of the ENS give rise to a range of disorders, termed enteric neuropathies and include conditions such as Hirschsprung’s disease. Little is known about the signalling that specifies early ENS progenitors. This has, thus far, limited progress in the generation of enteric neurons from human Pluripotent Stem Cells (hPSCs) that could provide a useful tool for disease modelling and regenerative medicine. We describe the efficient and accelerated generation of ENS progenitors from hPSCs, revealing that retinoic acid is critical for the acquisition of both vagal axial identity and early ENS progenitor specification. These ENS progenitors generate enteric neurons *in vitro* and following *in vivo* transplantation, achieving long-term colonisation of the ENS in adult mice. Thus, hPSC-derived ENS progenitors may provide the basis for cell therapy for defects in the ENS.

## Introduction

The enteric nervous system (ENS) is the largest branch of the peripheral nervous system and consists of an extensive network of neurons and glia controlling critical intestinal functions such as motility, fluid exchange, gastric acid/hormone secretion and blood flow (reviewed in (Furness, 2012; Sasselli et al., 2012). In amniote embryos, the ENS is derived predominantly from the vagal neural crest (NC), a multipotent cell population that is specified at the neural plate border between the presumptive neural and non-neural ectoderm between somites 1-7. Vagal NC also contributes to structures in various other organs such as the heart, thymus and lungs (Le Douarin et al., 2004; Hutchins et al., 2018; Simkin et al., 2018; Espinosa-Medina et al., 2017). After delaminating from the dorsal neural tube, vagal NC cells migrate first in a ventro-medial direction and enter the foregut. Following entry into the gut, enteric neural progenitors colonise the entire developing gut in a rostro-caudal direction. A number of studies have provided valuable insights into the determinants of ENS progenitor migration, proliferation and differentiation. These include the RET-GDNF (Heanue and Pachnis, 2008; Durbec, Marcos-Gutierrez, et al., 1996; Durbec, Larsson-Blomberg, et al., 1996) and Endothelin3-EDNRB (Baynash et al., 1994; Hosoda et al., 1994) signalling pathways as well as the transcription factors *SOX10*, *PHOX2B* and *ASCL1* (Elworthy et al., 2005; Bondurand et al., 2006; Memic et al., 2016). However, the signals that shape an early ENS identity within vagal NC precursors remain less well-defined.

The axial identity of vagal NC cells is characterized by the expression of members of the HOX gene paralogous groups (PGs) 3-5 (Parker and Krumlauf, 2017; Kam and Lui, 2015; Diman et al., 2011; Fu et al., 2003) and is patterned mainly by the action of somite-derived retinoic acid (RA) signalling, which acts by “posteriorising” cranial HOX-negative NC progenitors (Stuhlmiller and García-Castro, 2012; Frith et al., 2018; Ishikawa and Ito, 2009). Studies on *Xenopus* embryos have also indicated an earlier role for RA in neural crest induction (Villanueva et al., 2002; Bang et al., 1997). Further, gain- and loss-of-function studies in embryos have pointed to a critical role for RA in the specification of downstream vagal NC derivatives (Niederreither et al., 2001; Niederreither et al., 2003; Robrini et al., 2016). This role is especially prevalent in the development of the ENS where RA signalling components have been shown to control ENS progenitor migration and proliferation (Uribe et al., 2018) (Niederreither et al., 2003).

Human pluripotent stem cell (hPSCs) differentiation offers an attractive approach for dissecting the molecular and signalling basis of early developmental cell fate decisions. To date, a few studies have described the *in vitro* generation of ENS progenitors and enteric neurons from PSCs providing promising indications that these cell populations can be effectively utilised for the modelling and treatment of aganglionic gut conditions such as Hirschsprung disease (HSCR) (Barber et al., 2019; Fattahi et al., 2016; Workman et al., 2016; Kawaguchi et al., 2010; Schlieve et al., 2017; Lai et al., 2017; Li et al., 2016). These protocols rely on dual TGFβ/BMP inhibition to initially induce an anterior neuroectodermal intermediate that is subsequently converted into NC through WNT and BMP signalling while a vagal axial identity is induced through RA supplementation to eventually yield ENS progenitors after 10-15 days in culture (Fattahi et al., 2016; Lau et al., 2019; Workman et al., 2016). However, there is mounting evidence, both *in vitro* and *in vivo*, that the neural crest is in fact specified directly from cells of a pre-gastrulation identity through intermediate levels of BMP and early activation of the WNT signalling pathway (Basch et al., 2006; Hackland et al., 2017; Prasad et al., 2019). This suggests a more direct route of enteric neural crest induction is possible. Furthermore, the precise timing and concentration of RA signalling that controls the positional identity of vagal neural crest cells has not been clearly defined and it is not yet clear whether RA imparts an early enteric neural identity in hPSC-derived vagal neural crest or acts solely as a positional identity specifier.

We recently described a robust protocol for the efficient production of putative neural crest cells from hPSCs, (Hackland et al., 2017; Frith et al., 2018) that can acquire a vagal axial identity by exposure to RA (Frith et al., 2018). We have now utilised this *in vitro* differentiation system to thoroughly investigate the role of RA in both NC posteriorisation and ENS identity specification. We show that RA acts in a dose-dependent manner on pre-specified NC precursors, rather than pluripotent or neurectodermal cells, to induce expression of *HOX* genes indicative of a vagal character. This process appears to take place in parallel with the induction of early ENS progenitor markers. Crucially, we demonstrate that this effect of RA can be exploited to direct the accelerated production of ENS progenitors (within 6 days of differentiation), which are capable of generating enteric neurons and glia *in vitro* and which have the ability to colonise the ENS of adult mice following long-term transplantation. Our findings provide an efficient platform for the *in vitro* modelling of human ENS development and enteric neuropathies, as well as the development of cell therapy-based approaches for the treatment of such conditions.

## RESULTS

### The timing of retinoic acid signalling affects neural crest specification *in vitro*

We have previously shown that RA treatment of cranial NC precursors induces a vagal axial identity as defined by expression of HOX PG members 1-5 (Frith et al 2018). To define precisely the developmental time window during which RA exerts its action as a posteriorising signal without perturbing NC specification, we exposed differentiating hPSCs to 1μM all-*trans* RA at different stages of our NC differentiation protocol (Figure 1A). Induction of the NC markers p75 and *SOX10*, was assessed by flow cytometry following antibody staining in a SOX10-GFP reporter hPSC line (Chambers et al., 2012). Adding RA at day 0 of differentiation (the day of plating) did not result in any SOX10:GFP+/p75+ cells detected at day 5, whereas addition of RA at later time points (day 3 or 4 of differentiation), was compatible with the production of considerable numbers of SOX10:GFP+/p75+ cells (Figure 1B, C). Immunostaining for SOX10 expression in two additional independent hPSC lines (H7 and MasterShef7) confirmed the same temporal effect of RA addition on NC differentiation from hPSCs (Figure S1). These data suggest that RA signalling perturbs NC induction during the early stages of hPSC differentiation. Our findings also indicate that RA exerts its effects exclusively on cells committed to a NC fate rather than earlier ectodermal precursors or undifferentiated hPSCs.

**Figure 1:**
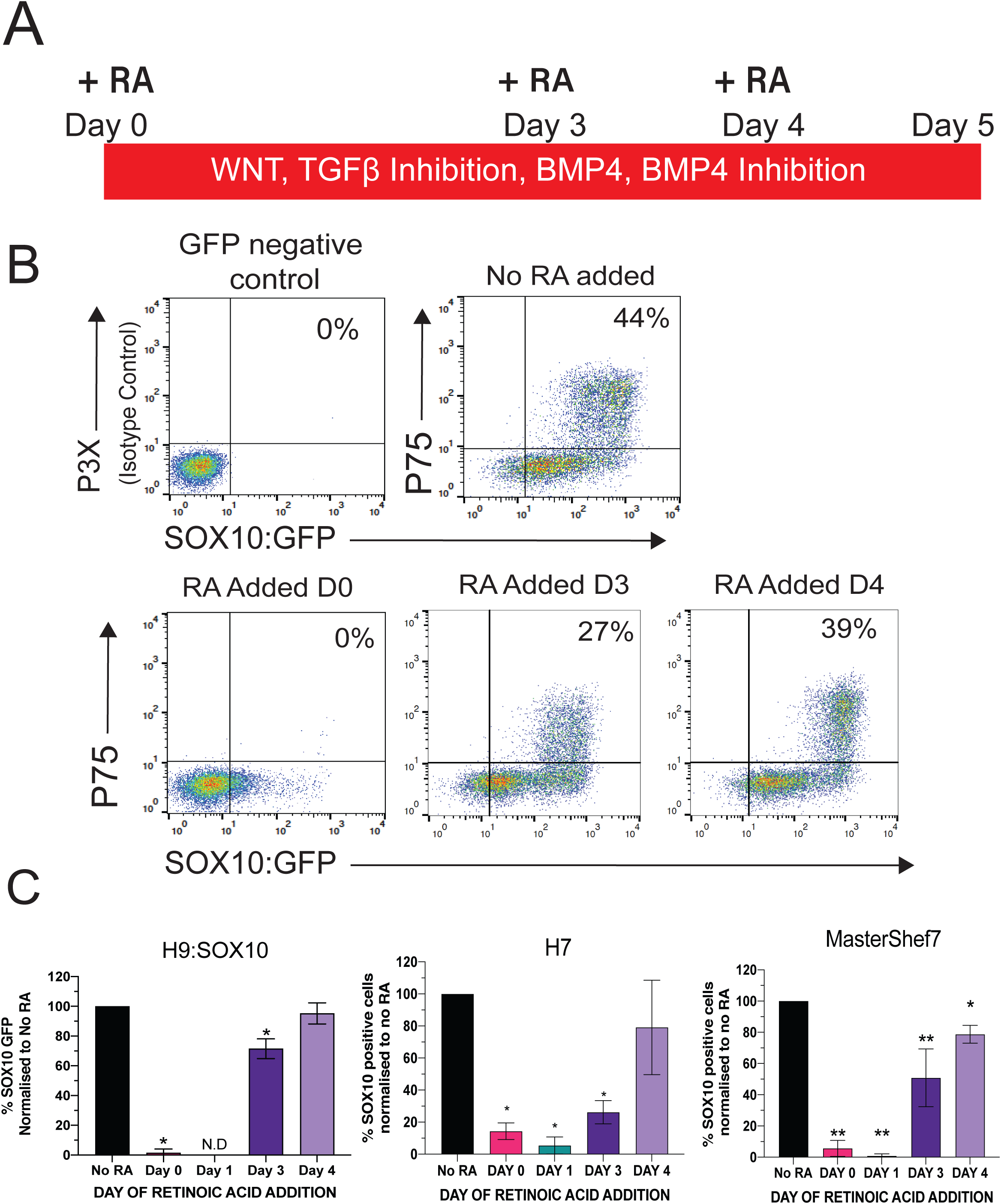
Retinoic Acid affects neural crest specification in a time-dependent manner. **(A)** Schematic showing the neural crest differentiation protocol and timepoints corresponding to addition of all-trans retinoic acid (RA). **(B)** FACs plots after 5 days of differentiation showing representative SOX10:GFP and p75 expression after RA addition at indicated timepoints during neural crest differentiation. Gates were set based on the negative control from a GFP negative line stained with P3X as a staining control. **(C)** Quantifications of the percentage of cells that display SOX10 expression in three different lines following FACs or immunofluorescence analysis. Graphs show percentages of SOX10+ cells normalised to a control condition where no RA was added. Bars state the mean ± standard deviation of 4 biological repeats for SOX10:GFP hPSCs and 3 biological repeats for H7 & MasterShef7. * P<0.05 ** P<0.01 One-way-Anova.

### Retinoic acid induces both vagal and enteric neural progenitor identities in a dose-dependent manner

RA has been shown to induce *HOX* gene expression in a dose-dependent manner *in vitro* (Okada et al., 2004; Simeone A, 1990) and *in vivo* (Papalopulu et al., 1991; Shimozono et al., 2013). To examine how the levels of RA signalling shape the acquisition of axial identities in hPSC-derived NC cells, we treated day 4 HOX-negative cranial NC precursors with varying concentrations of RA ranging from 10^−9^M (1nM) to 10^−6^M (1μM). This was followed by examination of the expression of various HOX genes as well as NC and ENS progenitor markers (Figure 2). We found that expression of *HOXB1* and *HOXB2*, was induced by RA at all concentrations in a dose-dependent manner (Figure. 2B, Figure. S2). In contrast, HOX genes marking vagal NC (HOXB4, B5 and B7) were only induced when higher concentrations of RA were employed (Figure. 2B). These data are consistent with previous findings showing that higher concentrations of RA induce more caudal identities (Okada et al., 2004; Simeone A, 1990). No expression of *HOXC9* was observed with any RA concentration, in line with other reports demonstrating that a trunk axial identity is mediated by WNT and FGF signalling (Frith et al., 2018; Frith and Tsakiridis, 2019; Abu-Bonsrah et al., 2018; Bel-Vialar et al., 2002; Lippmann et al., 2015; Mazzoni et al., 2013; Metzis et al., 2018; Denham et al., 2015; Hackland et al., 2019).

**Figure 2:**
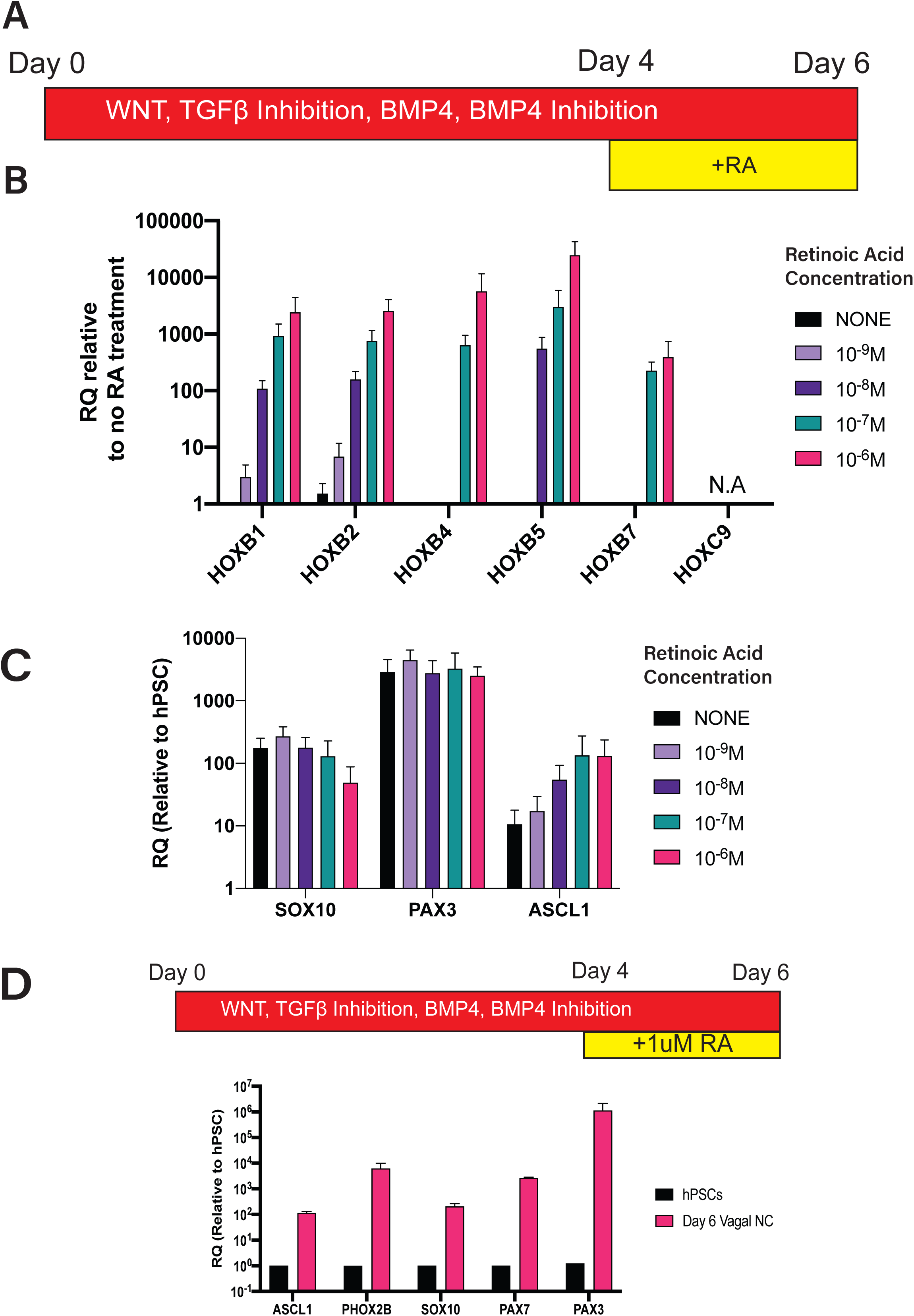
Retinoic induces a vagal axial and an early ENS progenitor identity in a dose dependent fashion. **(A)** Schematic showing the differentiation protocol with RA addition at day 4. **(B)** qPCR plots showing the induction of *HOX* genes in day 6 neural crest cells after exposure to different concentrations of retinoic acid. Data presented are relative quantities compared to *HOX* negative cells that were not treated with retinoic acid. Bars represent mean + standard deviation from 3 biological repeats of SOX10:GFP hPSCs. **(C)** qPCR plot showing the expression of the neural crest markers *SOX10*, *PAX3* and the enteric neural precursor marker *ASCL1* in day 6 cells following 2 days exposure to different concentrations of retinoic acid. Bars represent the mean + standard deviation of 3 biological repeats of SOX10:GFP hPSCs. **(D)** qPCR plots showing the expression of indicated neural crest and enteric neural precursor markers in day 6 cells that have been subjected to two days exposure of 1μM retinoic acid.

Expression of the NC markers *SOX10, PAX7* and *PAX3* was unaffected by the levels of RA (Figure 2C, Figure S2) confirming our previous observation that NC induction is not dependent on exogenous RA signalling (Figure 1). The highest concentrations of RA also triggered the initiation of an ENS progenitor transcriptional profile as defined by the expression of *ASCL1* and *PHOX2B* that mark migrating ENS progenitors (Blaugrund et al., 1996; Lo et al., 1991; Elworthy et al., 2005). Collectively, these results indicate that acquisition of a vagal axial identity and ENS progenitor specification in NC progenitors are tightly coupled events that are dependent on RA signalling.

### RA-induced vagal NC/ENS progenitors generate putative enteric neurons *in vitro*

To test whether day 6 RA-treated vagal NC cells treated with 1µM RA possess ENS progenitor potential we tested their ability to form enteric neurons *in vitro*. Day 6 RA-treated vagal NC cells were first cultured in the presence of WNT and FGF signals in non-adherent conditions to generate spheres (Figure 3A), as described previously (Fattahi et al., 2016). Flow cytometry and fluorescence microscopy analysis showed that these spheres retained SOX10:GFP expression and immunoreactivity to the NC markers p75 and CD49d (Figure 3B, C). Retention of an ENS progenitor identity in non-adherent culture conditions was also indicated by the sustained expression of ENS precursor markers *SOX10*, *PAX3*, *PAX7* and *ASCL1* (Figure 3D). Spheres were re-plated in conditions containing GDNF, ascorbic acid and NOTCH signalling inhibition, which promotes enteric neuron differentiation (Fattahi et al., 2016; Okamura and Saga, 2008; Theocharatos et al., 2013). One week following plating of spheres, we observed the emergence of cells with a neuronal morphology that expressed the enteric neuronal markers TUJ1, RET, TRKC and PERIPHERIN (Figure 3F). Similar results were obtained with two additional independent hPSC lines (Figure S3). Markers of both enteric neurons and glia were detected by quantitative real time PCR (qPCR) in day 22 cultures (Figure 3G). Furthermore, *ChAT*, 5-*HT, TH* and *ASCL1* expression further confirmed the presence of early enteric neurons in the cultures (Figure 3G). Transcripts for the early glial markers *SOX10* and *S100β* were also detected in day 22 cultures, but expression of *GFAP*, that is characteristic of more mature enteric glia, was not observed (**Data not shown)**. Together these observations suggest that day 6 RA-induced NC cells can give rise to enteric neurons and glia *in vitro*.

**Figure 3:**
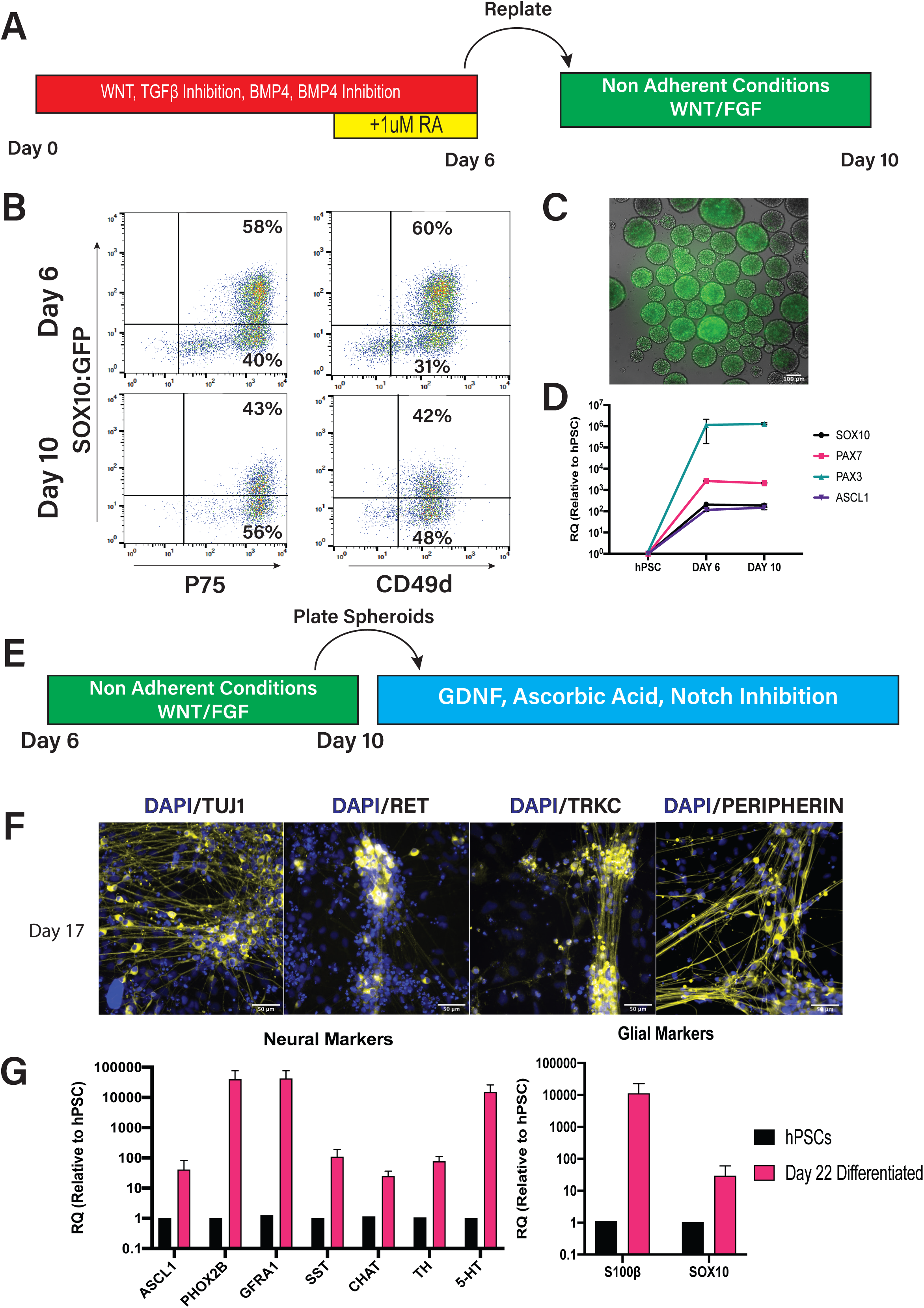
Day 6 putative enteric neural precursors can generate putative enteric neurons *in vitro*. **(A)** Schematic to depict the culture of day 6 cells in non-adherent conditions. **(B)** FACs plots showing the retention of SOX10:GFP and p75/CD49d co-expression from day 6 to day 10 after culture in non-adherent conditions with WNT/FGF. **(C)** Representative photomicrograph of neural crest spheres at day 8 showing SOX10:GFP positive cells forming spheres. **(D)** qPCR graph showing that the expression of the neural crest markers SO*X10, PAX7, PAX3* and the enteric neural precursor marker *ASCL1* is retained between day 6 and day 10 following culture in non-adherent conditions. Bars represent mean±standard deviation of 3 biological repeats. **(E)** Schematic depicting the conditions to generate enteric neurons from day 10 spheres following plating. **(F)** Immunofluorescence analysis shows the presence of cells that are positive for TUJ1, RET, TRKC and PERIPHERIN at day 17 of differentiation. Scale bars are 50μm. **(G)** qPCR analysis shows the induction of markers for early enteric neurons and early glial progenitors at day 22 of differentiation. Expression is shown as relative quantity compared to hPSCs. Bars represent mean + standard deviation of 3 biological repeats. All data shown have been obtained using SOX10:GFP hPSCs.

### RA-induced vagal NC/ENS progenitors colonise the adult mouse ENS *in vivo*

To assess the developmental potential of hPSC-derived vagal neural crest/ENS progenitors *in vivo*, we performed transplantations into the caecum of adult immunodeficient Rag2^−/−^;γc^−/-^;C5^−/−^ mice. To track cells post transplantation, we used the human induced pluripotent stem (iPS) cell line SFCi55-ZsGr that contains a constitutive ZsGreen fluorescent reporter in the *AAVS* safe harbour locus (Lopez-Yrigoyen et al., 2018). ZsGreen+ iPS cells were differentiated toward vagal NC/ENS progenitors as described above (Figures 2 & 3) and spheres were generated from sorted p75^++^ cells (Figure 4A, B). The cells were then transplanted to the serosal aspect of the caecum in adult (4-8 week old) immunodeficient Rag2^−/−^;γc^−/−^;C5^−/−^ mice and analysed for integration and differentiation at timed intervals. At 2 weeks post-transplantation ZsGreen+ cells were observed at the serosal aspect both within the caecum and proximal colon. ZsGreen+ cells expressing the neuronal marker TUJ1 were also observed (Figure 4C; left; Arrowheads). Such ZsGreen+/TuJ1+ cells were found to form filamentous and interconnecting projections along the serosal surface at this timepoint. At 4 weeks post-transplantation ZsGreen+ cells were again observed on the serosal surface at the presumptive site of transplantation. Importantly, ZsGreen+ cells were also found within the *tunica muscularis* at the level of the myenteric plexus. Within the *tunica muscularis* ZsGreen+ cells were found to co-express the neuronal marker TUJ1 both within myenteric ganglia-like structures and as intramuscular neurons (Figure 4C; right). Additionally, ZsGreen+ cells, which were also positive for the glial marker GFAP and located at the level of the myenteric plexus were also detected (Figure 4C; right) suggesting that hPSC-derived ENS progenitors have the potential to differentiate to the main enteric cell types after transplantation *in vivo*. Moreover, 3 months after transplantation ZsGreen+ cells could be identified across the gut wall both within individual myenteric ganglia (Figure 4D; left) and within the submucosa, surrounding cryptal structures (Figure 4D; right). At this timepoint, we observed the presence of ZsGreen+ cells which had differentiated into major enteric neuronal subtypes defined by the expression of either neuronal nitric oxide synthase (nNOS; Figure 4D; left) or vesicular acetylcholine transporter (vAChT; Figure 4D; right). Together these results suggest that hPSC-derived ENS progenitors can integrate into recipient gut tissue where they are maintained in the long-term and have the ability to differentiate to multiple neuronal subtypes and glia.

**Figure 4:**
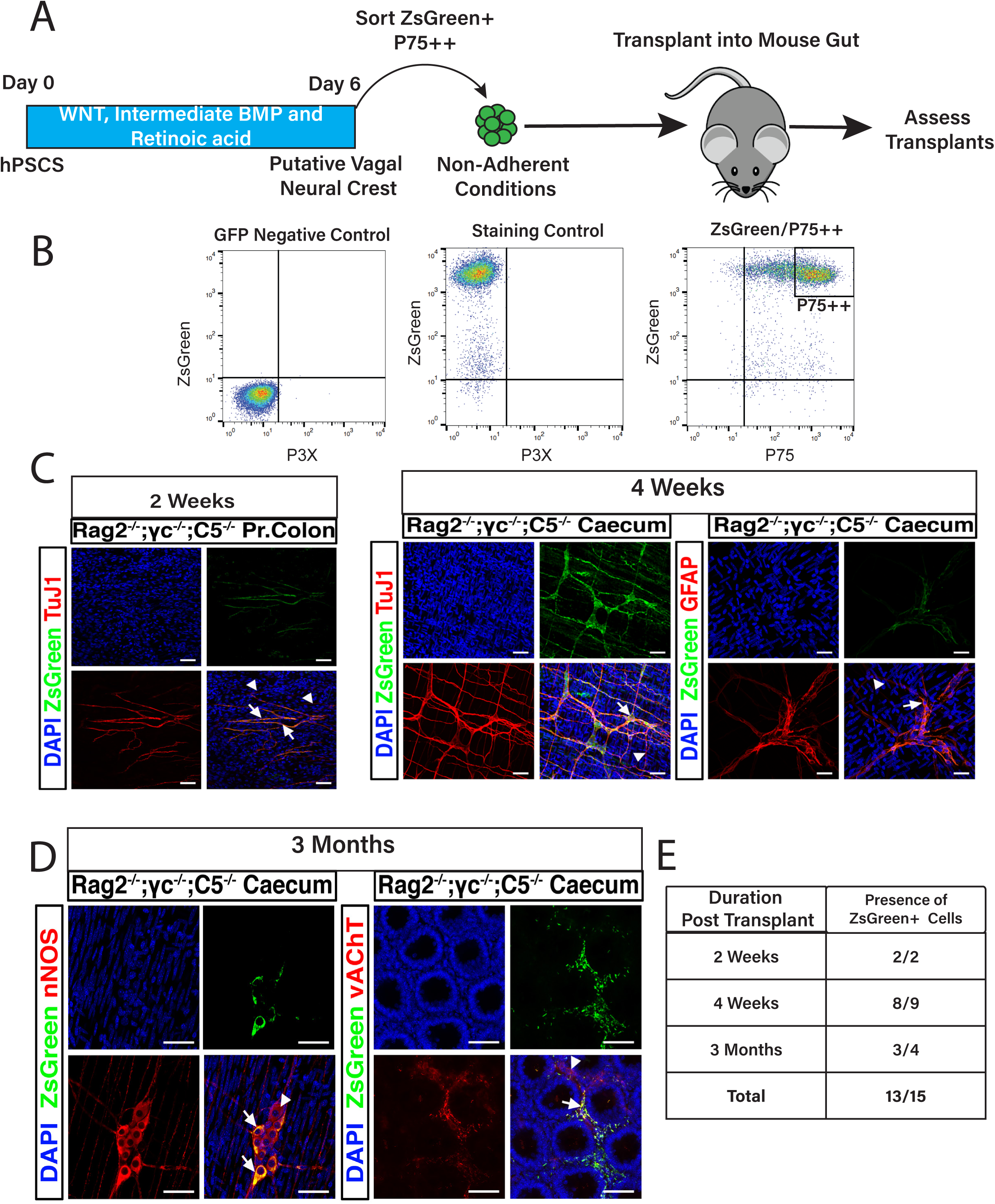
hPSC derived enteric neuronal precursors integrate into the mouse ENS after transplantation. **(A)** Schematic depicting the experimental procedures for transplantation of hPSC-derived ENS progenitors **(B)** Sorting strategy to isolate ZsGreen+/p75++ labelled putative ENS progenitors following *in vitro* differentiation. P3X antibody is control for antibody staining. **(C)** Wholemount images of gut tissue corresponding to the indicated regions obtained from immunodeficient mice at 2 weeks and 4 weeks post-transplantation and showing the presence of ZsGreen+ cells that are positive for the neuronal marker TUJ1 (Arrows) amongst endogenous TUJ1+ neurons (Arrowheads), and glial marker GFAP after immunostaining. Pr. Colon, proximal colon. **(D)** Images showing the differentiation of hPSC-derived ENS progenitors into different subtypes of enteric neurons including nNOS positive and vAChAT positive neurons in the caecum of Rag2^−/−^;γc^−/−^;C5^−/−^ mice at 3 months post-transplantation. Arrows are transplanted ZsGreen+ cells; Arrowheads are endogenous enteric neurons. **(E)** Table showing the numbers of mice in which ZsGreen+ cells were identified over the total number of transplanted mice analysed at indicated timepoints post-transplantation.

## Discussion

We have described an efficient differentiation system that can be employed in multiple hPSC lines, based on the use of retinoic acid to drive the concomitant induction of both a vagal and an ENS progenitor identity in cranial neural crest progenitor cells. The generation of early ENS progenitors after just 6 days, is quicker than other previously published protocols that describe the production of similar cell populations from hPSCs (Fattahi et al., 2016; Workman et al., 2016; Barber et al., 2019). The common feature of these protocols is the induction of an anterior neuroectodermal intermediate by dual-SMAD inhibition to produce NC that then becomes “posteriorised” into a vagal axial identity by RA to give rise to ENS progenitors after 10-15 days. A likely reason for the accelerated production of similar progenitors in the protocol we describe is that we employ a strategy that generates an NC precursor population from hPSCs through WNT and intermediate levels of BMP signalling, which then acquires a vagal/ENS identity through the action of RA. This is in line with our previous findings (Frith et al., 2018), as well as with other studies (Basch et al., 2006; Leung et al., 2016; Prasad et al., 2019) suggesting that NC specification occurs independently of a neurectodermal precursor intermediate. Our protocol does not include any serum replacement and is fully defined to reduce variability within components of the differentiation media. We also use ‘Top-Down Inhibition’ to control BMP signalling to reduce further variability and improve robustness across independent hPSC lines (Hackland et al., 2017).

RA also appears to drive specification of early ENS progenitors as indicated by expression of *SOX10*, *ASCL1* and *p75* (Figure 2), consistent with previous studies that reveal a critical role for RA signalling in promoting ENS progenitor migration, proliferation and differentiation (Niederreither et al., 2003; Uribe et al., 2018; Simkin et al., 2013). Further, ENS progenitor identity is acquired alongside a vagal axial identity during neural crest specification, in a manner that is dependent on the dose of RA. This finding suggests that the gene regulatory networks that control axial identity and cell fate are inter-linked and inter-dependent. RA signalling may control these processes through the induction of vagal level HOX family members such as *HOXB3* and *HOXB5* as well as their TALE family co-factors, such as the *MEIS* family of genes, which have been reported to regulate downstream ENS development (Chan et al., 2005; Kam et al., 2014; Kam and Lui, 2015; Uribe et al., 2018; Uribe and Bronner, 2015) by acting upstream of *Ret* (Zhu et al., 2014) and preventing apoptosis (Kam et al., 2014).

Critically, our differentiation strategy rapidly yields (by day 6 of differentiation) a promising well-defined cell population that can efficiently generate enteric neurons and glia in vitro (Figure 3), and so may provide a potential cellular donor for the treatment of enteric neuropathies. In contrast to previous *in vivo* studies utilising hPSC-derived ENS progenitors, we chose to transplant our ENS progenitors into the gut of immunodeficient Rag2^−/−^;γc^−/−^;C5^−/−^ mice. This approach eliminates the requirement for chemical immunosuppression and allows long-term study of donor cell survival and integration within a “normal” host ENS microenvironment. Crucially, we found that the hPSC-derived neurons were present within endogenous ENS ganglia of adult mice up to 3 months post-transplantation, expressing the same characteristic markers (nNOS and vAChT positive neurons) as the mouse neurons in their host environment (Figure 4). Further, the transplanted human cells populated both the myenteric and submucosal plexuses of the gut, demonstrating that they are able to migrate extensively within the gut wall and form neuronal networks even though the host ENS remained intact (Figure 4). Recent studies have demonstrated the potential of cellular transplantation for the treatment of enteric neuropathies. Importantly, both postnatally-derived human and murine endogenous enteric neural stem cells (ENSC) have been used for *in vivo* applications, in mice, demonstrating functional integration (J.E. Cooper et al., 2017; J.E. Cooper et al., 2016; Stamp et al., 2017), and functional rescue of an enteric neuropathy (McCann et al., 2017). Similarly, transplanted hPSC-derived ENS progenitors generated through dual-SMAD inhibition have been shown to integrate and migrate extensively within a mouse model of Hirschsprung disease leading to increased survival (Fattahi et al., 2016). Our work here extends and complements these studies providing further evidence in support of the use of hPSCs as a promising platform for the development of cell therapies to treat ENS dysfunction.

## Methods and Materials

### hPSC culture

The human pluripotent stem cell lines H7 (WA07), H9 (WA09) (Thomson, 1998), H9:SOX10 (Chambers et al., 2012), Mastershef7 (Gouti et al., 2014), and SFCi55-ZsGr (Lopez-Yrigoyen et al., 2018) were grown in mTESR (Stem Cell Technologies # 85850) on 1:100 dilution of Geltrex (ThermoFisher A1413202) in DMEM/F12 (Sigma D6421). Cells were passaged at 80-90% confluency using ReLeSR (Stem Cell Technologies Catalog # 05873). Cells were incubated at 37°C in 5%CO_2_.

### Directed Differentiation

For vagal neural crest differentiation, we use a previously described protocol (Frith et al., 2018). hPSCs at approximately 80% confluency were detached using Accutase (Sigma-Aldrich A6964) for 10 minutes at 37°C to generate single cells. Cells were counted manually and plated at 50,000 cells/cm^2^ on Geltrex coated plates (ThermoFisher A1413202). Neural Crest differentiation media is comprised of DMEM/F12 (Sigma-Aldrich), supplemented with 1x N2 (ThermoFisher 17502048), NEAA (ThermoFisher 11140050), Glutamax (ThermoFisher 35050061), 1μM CHIR99021 (Tocris 4423), 2μM SB431542 (Tocris 1614/1), 1μM DMH-1 (Tocris 4126/10), 20ng/ml BMP4 (ThermoFisher PHC9533). All-Trans Retinoic Acid (Sigma R2625) was diluted in DMSO. 10μM Y-27632 dihydrochloride (Tocris 1254/1) was added at day 0 until day 2 to assist attachment. For all vagal neural crest induction all-trans Retinoic acid was added at a final concentration of 1μM on day 4 unless specified in the results. Media was changed every other day until day 5/6.

Spheres were generated as previously described (Fattahi et al., 2016). Day 6 cells were treated with accutase to form a single cell suspension and replated in a media containing a 1:1 mix of DMEM/F12 (Sigma) with Neurobasal (ThermoFisher 21103049) supplemented with 1x N2, 1x B27, 1x NEAA, 1x Glutamax, 3μM CHIR99021, 10ng/ml FGF2 (R&D systems 233-FB/CF). Sphere media supplemented 10μM of Y-27632 dihydrochloride (Tocris) to ensure sphere formation and left until day 10. One well of a 6 well plate was plated into one well of an Ultra-Low Attachment 6 well plate (Corning 3471) or 6 well plates with a coating of 1% w/v agarose.

For enteric neuronal differentiation, day 10 spheres were plated onto Geltrex coated plates in BrainPhys (Stem Cell Technologies 05790) supplemented with 1x N2, 1x B27 (ThermoFisher 17504044), 100μM Ascorbic Acid (Sigma A8960), 10ng/ml GDNF (Peprotech 450-10) and 10μM DAPT (Sigma D5942). Media was changed every other day and once a week supplemented with Vitronectin (ThermoFisher A14700)

### RNA extraction, cDNA synthesis & qPCR

RNA was extracted using a Total RNA purification plus kit (Norgen BioTek #48300) per manufacturer’s instructions. RNA concentration was measured using a nanodrop (ThermoFisher). RNA was stored at −80°C. cDNA was synthesised using the High-Capacity cDNA Reverse Transcription kit (ThermoFisher 4368813) and stored at - 20°C.

qPCR was performed on QuantStudio 12K Flex thermocycler (Applied Biosystems). CT values were calculated against GAPDH for each sample. Relative quantities calculated using the −2^^CT method.

### Immunofluorescence

Cells were fixed with 4% PFA for 10 minutes at room temperature and washed 3 times with 1x PBS (no Mg^2+^/ Ca^2+^). Cells were permeabilised and blocked with 1x PBS (no Mg^2+^/ Ca^2+^) supplemented with 10% FCS, 0.1% BSA and 0.3% Triton-X 100 for 1 hour at room temperature. Primary antibodies were diluted in permeabilisation buffer and incubated at 4°C overnight. Secondary antibodies were diluted in permeabilization buffer and stained in the dark at 4°C for one hour. Nuclei were counterstained with Hoechst 33342 (ThermoFisher H3570). Images were taken on an InCell Analyser 2500 (GE Healthcare).

### Image Analysis

Images were quantified using custom made pipelines on CellProfiler 2.2 (Carpenter et al., 2006). Individual nuclei were identified by Hoechst staining and the fluorescence intensity following staining was measured and related to nuclei. Positive staining was scored based on having greater fluorescence intensity values than a threshold value from a secondary only staining control for each biological repeat.

### Flow Cytometry

Flow cytometry was carried out as described in (Frith et al., 2018). In short, a single cell suspension was generated using Accutase as described above. Cells were pelleted and resuspended in FACs buffer (DMEM/10% v/v FCS) at 1×10^6^ cells/ml. Gating for positive cells was based on a negative control consisting of cells not carrying a reporter or cells stained with P3X, an antibody from the parent myeloma (KOHLER and MILSTEIN, 1975; Hackland et al., 2017).

#### Animals

Animals were maintained, and experiments were performed, in accordance with the UK Animals (Scientific Procedures) Act 1986 under license from the Home Office (P0336FFB0) and approved by the University College London Biological Services Ethical Review Process. Animal husbandry at UCL Biological Services was in accordance with the UK Home Office Certificate of Designation.

Rag2^−/−^;γc^−/−^;C5^−/−^ mice, which lack innate immunity and are deficient in all lymphocytes (R.N. Cooper et al., 2003; Silva-Barbosa et al., 2005),were used as recipients for all hPSC-derived vagal neural crest/ENS progenitor transplantations.

### *In vivo* Cell Transplantation

ZsGreen+ spheres were transplanted to the caecum of 4-8 week-old immunodeficient Rag2^−/−^;γc^−/−^;C5^−/−^ mice, via laparotomy under isofluorane anesthetic. Briefly, the caecum was exposed and ZsGreen+ spheres, containing 1 million cells each, were subsequently transplanted to the serosal aspect of the caecum by mouth pipette, using a pulled glass micropipette. Each transplanted tissue typically received 3 ZsGreen+ spheres which were manipulated on the surface of the caecum with the bevel of a 30G needle to ensure correct positioning. Transplanted Rag2^−/−^;γc^−/−^;C5^−/−^ mice were typically maintained for up to 3 months post-transplantation before sacrifice and removal of the caecum and proximal colon for analysis.

### Wholemount Gut Immunohistochemistry

Wholemount immunohistochemistry was performed on transplanted caecal and proximal colon segments after cervical dislocation and excision. Tissues were fixed in ice cold 4% PFA for 45 min at 22°C. After fixation, tissues were washed for 24h in 1x PBS at 4°C. Cells were permeabilised and blocked with 1x PBS supplemented with 1% Triton X-100 and 10% sheep serum. Primary antibodies were diluted in permeabilisation buffer and incubated at 4°C for 48h. Secondary antibodies were diluted in permeabilization buffer and stained in the dark for one hour at 22°C. Nuclei were counterstained with DAPI (Sigma). Before mounting, tissues were washed thoroughly in 1x PBS for 2h at 22 °C. Tissues were examined using a LSM710 Meta confocal microscope (Zeiss). Confocal micrographs of whole mounts were digital composites of the Z-series of scans (0.5-1μm optical sections, 10–50μm thick).

### Antibodies

|  | **Antibody** | **Species** | **Source** | **Dilution** |
| --- | --- | --- | --- | --- |
| In Vitro | SOX10 | Rabbit | Cell Signalling Technology (D5V9L) #89356 | 1:500 |
|  | RET | Rabbit | Abcam ab134100 | 1:1000 |
|  | TUJ1 | Mouse | Abcam ab78078 | 1:1000 |
|  | TRKC | Rabbit | Cell Signalling Technology (C44H5) #3376 | 1:1000 |
|  | PERIPHERIN | Rabbit | Millipore AB1530 | 1:100 |
|  | P3X | Mouse | In House myeloma P3X63Ag8 (Kohler & Milstein 1975) | 1:10 |
|  | p75 | Mouse | In House Hybridoma Clone ME20.4 (Ross et al., 1984) | 1:20 |
|  | CD49d | Mouse | BioLegend 304302 Clone 9F10 | 1:100 |
| Transplants | TUJ1 | Mouse | BioLegend MMS-435P | 1:500 |
|  | GFAP | Rabbit | Millipore AB5804 | 1:500 |
|  | nNOS | Rabbit | Invitrogen 61-7000 | 1:400 |
|  | vAChT | Goat | ThermoFisher Scientific OSH00003W | 1:200 |
|  | DAPI |  | Sigma D8417 | 1:1000 |

### Primers

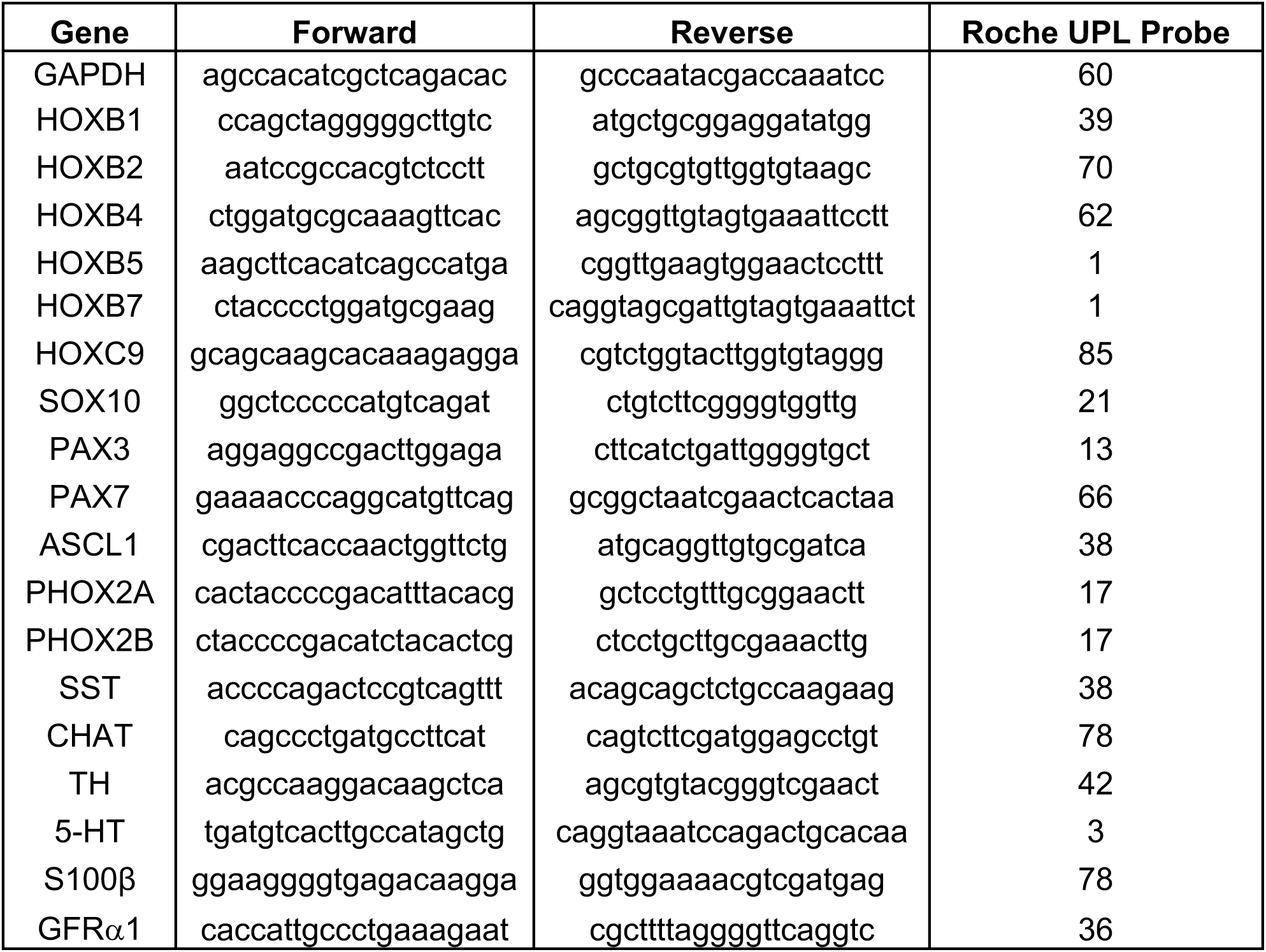

## Acknowledgements

This work was supported in part by grants from the Medical Research Council Confidence in Concept awarded to IB & PWA (MC_PC_14115), BBSRC (BB/P000444/1) awarded to AT, H2020-EU.1.2.2 (Grant agreement ID: 824070) awarded to AT & PWA. CM is supported by Guts UK (Derek Butler Fellowship). NT is supported by Great Ormond Street Hospital Children’s Charity (GOSHCC - V1258). This work was partially funded through a GOSHCC grant (W1018C) awarded to NT (Principal Investigator) and AJB (Co-Investigator).

The authors would like to acknowledge the NIHR Great Ormond Street Hospital Biomedical Research Centre which supports all research at Great Ormond Street Hospital NHS Foundation Trust and UCL Great Ormond Street Institute of Child Health. The views expressed are those of the author(s) and not necessarily those of the NHS, the NIHR or the Department of Health. The authors acknowledge the support of Prince Abdullah Ben Khalid Celiac Research Chair, College of Medicine, Vice-Deanship of the Research Chairs, King Saud University, Riyadh, Saudi Arabia.

## Author Contributions

TF, PWA, JOSH, CM, AJB, NT conceived the project. TF, CM designed, performed and analysed experiments with help from AT & AG.

IB, PWA, AT, NT, AJB and CM provided financial support. TF, AT, CM and PWA wrote the manuscript.

---

**Supplementary Figure 1:**
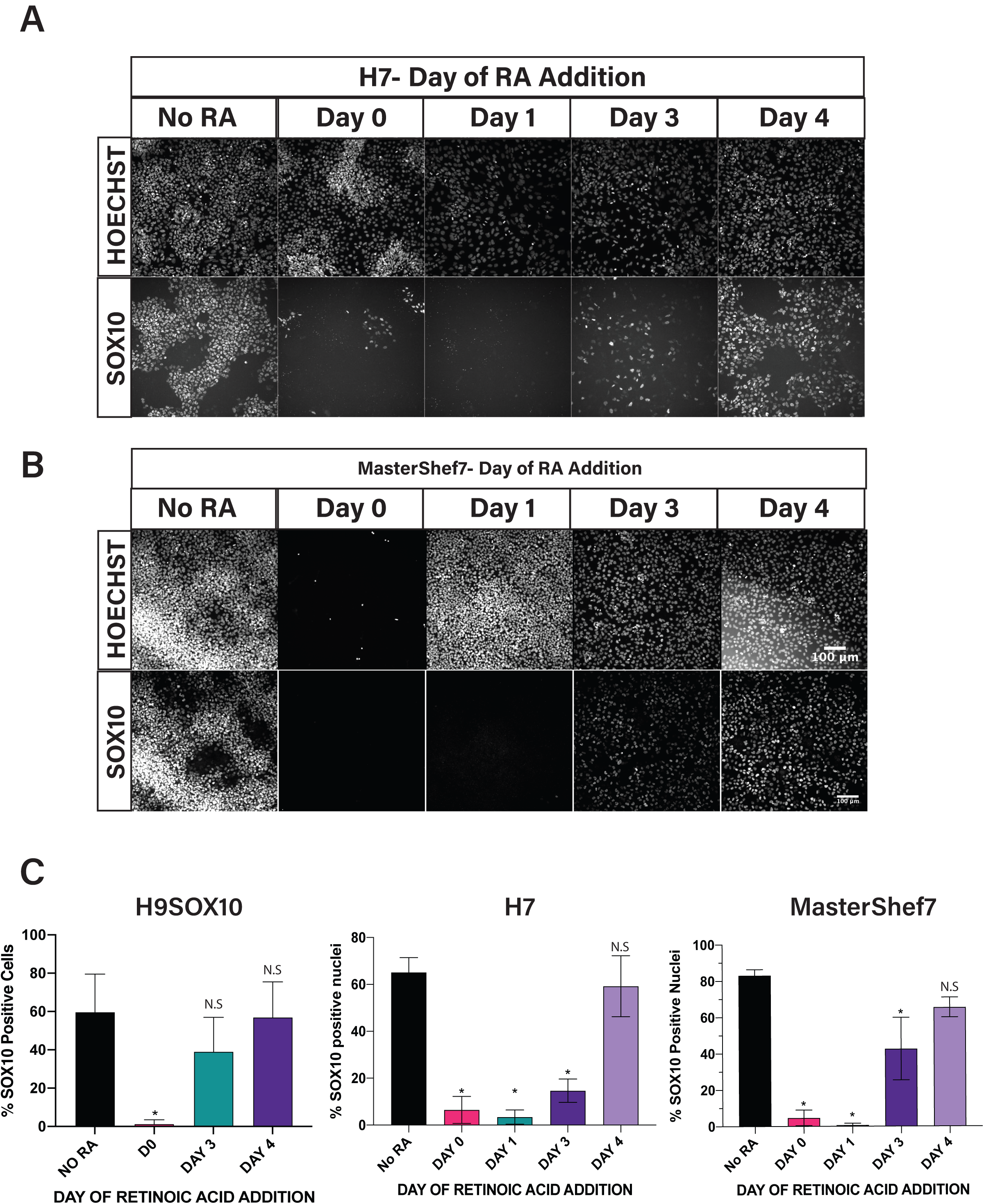
RA timing conserved across other hPSC lines. **(A-B)** Immunofluorescence images showing SOX10 expression at day 5 after RA was added at different times during neural crest differentiation of H7 **(A)** and MasterShef7 **(B)** hPSCs. **(C)** Quantification of the number of GFP positive cells from SOX10:GFP hPSCs (4 biological repeats) and SOX10 positive cells in H7 and MasterShef7 after addition of RA at different time points from 3 biological repeats. Bars are mean ± standard deviation. *P<0.05, N.S= not significant One-Way Anova compared to day 5 cells not treated with RA.

**Supplementary Figure 2:**
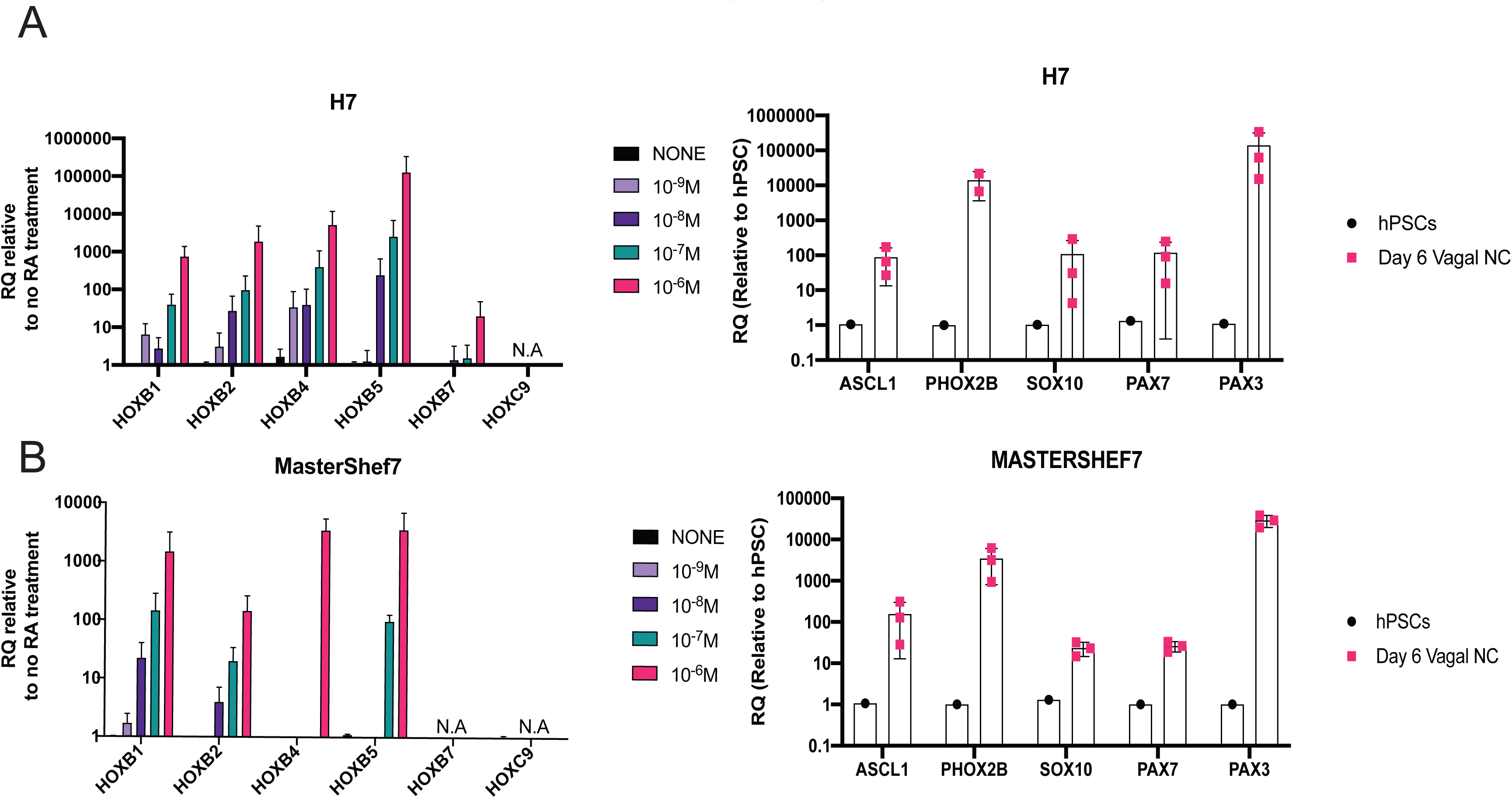
HOX gene induction and early enteric neural marker induction is dependent on the concentration of RA. qPCR analysis showing HOX gene and early ENS progenitor marker induction after 6 days of differentiation following RA exposure of H7 **(A)** and MasterShef7 **(B)** hPSCs

**Supplementary Figure 3:**
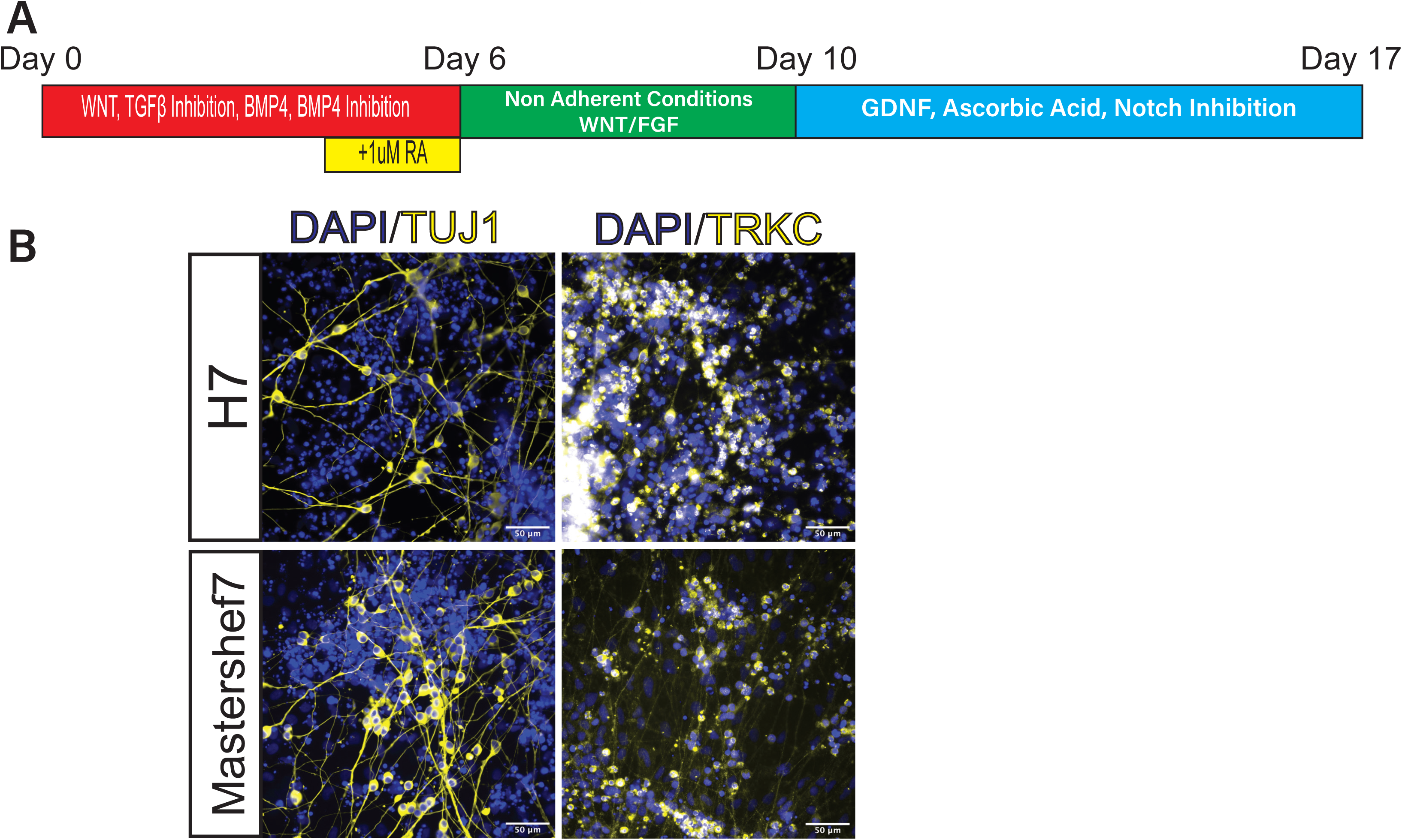
Generation of enteric neurons in 2 further independent hPSC lines. **(A)** Schematic of enteric neuron differentiation protocol. **(B)** Immunofluorescence showing the cells that are positive for TUJ1 and TRKC at day 17 of differentiation. Scale bars are 50μm.

